# Substrate sensing institutes sequential and asymmetric electron transfer in the nitrogenase-like DPOR complex

**DOI:** 10.1101/2020.03.31.018135

**Authors:** Elliot I. Corless, Brian Bennett, Edwin Antony

## Abstract

Dark-operative protochlorophyllide oxidoreductase (DPOR) catalyzes the reduction of protochlorophyllide (Pchlide) to chlorophyllide (Chlide), a key penultimate step in the biosynthesis of bacteriochlorophyll. DPOR shares structural homology with nitrogenase and is made of electron donor (BchL) and electron acceptor (BchNB) component proteins. ATP driven assembly of the BchL and BchNB proteins drives electron transfer and Pchlide reduction. BchNB is composed of two subunits each of BchN and BchB arranged as an α2ß2 heterotetramer. Here, we describe extensive allosteric communication between the two identical active sites in BchNB that drives sequential and asymmetric electron transfer. Pchlide binding and electron transfer activities in one half of the BchNB tetramer allosterically regulates activities in the other half. Pchlide binding is sensed and recognized *in trans* by an Asp274 from the opposing half and is positioned in the active site to likely serve as the initial proton donor. An Asp274 to Ala substituted DPOR binds to two Pchlide molecules in the BchNB complex but is unable to conformationally poise one Pchlide molecule. Thus, stalling Pchlide reduction in both active sites. The [4Fe-4S] cluster of the BchNB protein is pre-reduced and donates the first electron to Pchlide, a mechanism similar to the deficit-spending model observed in nitrogenase. In half-reactive DPOR complexes, incapacitating proton donation in one half generates a stalled intermediate and Pchlide reduction in both halves is abolished. The results showcase long-range allosteric communication and sequential ET in the two symmetric halves. The findings shed light on the functional advantages imparted by the oligomeric architecture found in many electron transfer enzymes.

## Introduction

Many oligomeric enzymes that transfer electrons for catalysis or substrate reduction have two identical active sites and their subunits are arranged with a head-to-head or head-to-tail symmetry. The electron acceptor component proteins of Nitrogenase and the nitrogenase-like class of enzymes such as the dark operative protochlorophyllide oxidoreductase (DPOR) and chlorophyllide oxidoreductase are arranged as α_2_β_2_ tetramers (Fig. 1a)(1–4). A similar architecture is also seen in the nitric oxide synthase(5) and ribonucleotide reductase family of enzymes(6). Given the evolutionary and functional significance of these enzymes, a mechanistic significance must exist behind such structural assemblies. In gymnosperms, the cyanobacteria *Prochlorococcus marinus* and all photosynthetic eubacteria, DPOR catalyzes the reduction of protochlorophyllide (Pchlide) to chlorophyllide (Chlide), the penultimate step in the biosynthesis of chlorophyll and bacteriochlorophyll(7).

**Figure 1.**
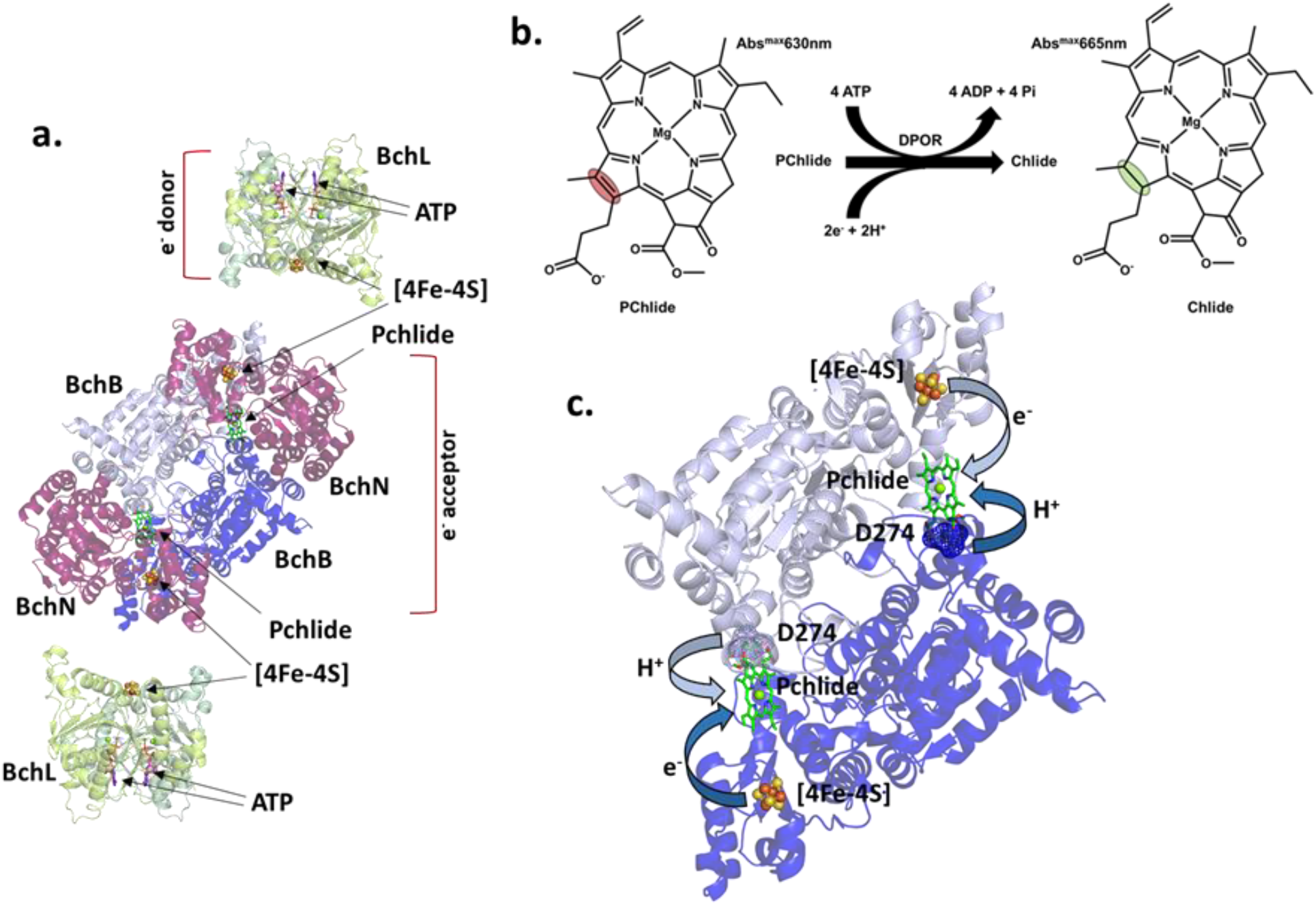
The α_2_β_2_ asymmetric structural arrangement of BchNB. **a.** Crystal structure of the electron donor (BchL) and electron acceptor (BchNB) protein components of DPOR (PDB:2YNM). The homodimeric BchL bound to ATP is shown. The BchNB tetrameric complex is shown with both BchN subunits in red, and the two BchB subunits are colored light and dark blue, respectively. Pchlide, metal clusters and ATP are shown as sticks. **b.** Schematic of Pchlide reduction to Chlide by DPOR. Two cycles of electron transfer from BchL to BchNB are required for reduction of the C17=C18 double bond (marked by colored ovals). **c.** Crystal structure of the two BchB subunits are shown to highlight the position of Asp274 relative to Pchlide and the [4Fe-4S] cluster. Electron transfer from the [4Fe-4S] cluster to Pchlide and proton donation from D274 are denoted by colored arrows.

DPOR consists of electron donor (BchL) and electron acceptor (BchN-BchB; BchNB) component proteins (Fig. 1a)(8, 9). BchL is a homodimer containing one [4Fe-4S] cluster ligated at the dimer interface by two cysteine residues per monomer and possesses one ATP binding site per monomer(10). BchNB is a α_2_β_2_ tetramer carrying one [4Fe-4S] cluster and substrate (Pchlide) binding site per half of the tetramer(3). ATP binding to BchL drives the assembly of the BchL and BchNB proteins and the transient assembly of this complex promotes electron transfer (ET)(8). The electron is transferred from the [4Fe-4S] cluster of BchL, to the [4Fe-4S] cluster of one half of BchNB and finally to the C17=C18 double bond of Pchlide. Two rounds of electron and proton transfer and are required to reduce Pchlide to Chlide (Fig. 1b). In DPOR, one proton required for reduction originates intrinsically from within the C17 propionate of Pchlide, and the second proton is donated in *trans* from an Asp274 of the opposing BchB subunit (Fig. 1c)(11, 12). Chlide is subsequently reduced by the structurally homologous Chlorophyllide Oxido-Reductase (COR) in the bacteriochlorophyll biosynthetic pathway(13, 14). These reductive steps fine tune the spectral and reactive properties of bacteriochlorophyll to be optimally suited for photosynthesis.

An overarching question about these enzymes is centered on their conserved architectural complexity: Why are these enzymes assembled as two functional halves, and how do they cooperatively function during substrate reduction? Since two rounds of ET (per half) are required for substrate reduction, multiple BchL binding and dissociation cycles occur at each BchNB half. The two BchL binding interfaces on BchNB are situated ~100Å apart(4). If their binding events are coordinated, long-range, inter-subunit allosteric communication is necessary. Here, we explored such fundamental mechanistic questions using DPOR. We show that the two halves indeed communicate, and that perturbation of substrate reduction activity at one half abolishes ET at the other, thus stalling the entire DPOR complex. Unique EPR signatures of the [4Fe-4S] cluster of the BchNB protein provide evidence that in the presence of BchL and ATP, BchNB can be reduced before binding of Pchlide, enabling the first ET to Pchlide in the absence of the BchL protein. This process is similar to the ‘deficit-spending’ mechanism observed in nitrogenase(15). Finally, we show that Pchlide binding to BchNB sets the functional asymmetry within the complex and the initial ET event is used as a sensory mechanism to recognize Pchlide and trap it in the active site. We propose that the α_2_β_2_ architecture in DPOR serves to correctly recognize and orient the incoming substrate. The recognition then triggers sequential ET which is coordinated through allosterically controlled conformational changes between the two halves of the BchNB complex. The findings explain key functional advantages of the oligomeric architecture found in many enzymes that catalyze electron transfer reactions.

## Results

### A half-reactive DPOR complex is defective for substrate reduction

To test the importance of the α_2_β_2_ architecture for function, we first generated a half-reactive version of DPOR and tested overall Pchlide reduction activity. BchN and BchB are expressed as separate open reading frames and form a constitutive tetramer. To generate a half-reactive BchNB complex, we used a dual-affinity tag approach where two BchB open reading frames were generated, coding for either a poly-histidine affinity tag or a Strep-tag affinity on the N-terminus. N-terminal positioning of either affinity tag does not affect Pchlide reduction activity as their rates of Pchlide reduction are similar (Fig. 2a, b). Reduction of Pchlide to Chlide results in a spectral shift with appearance of an absorption maxima at 666 nm. We note that placing affinity tags on the C-terminus of BchB perturbs Pchlide reduction activity (data not shown). These tagged-BchB constructs were co-expressed along with an untagged version of the BchN polypeptide. This strategy generated an ensemble of BchNB tetramers carrying either two poly-histidine tagged BchB, two step-tagged BchB, or one of each in the context of a BchNB tetramer (Fig. 2c). This dual-tagged approach enabled us to sequentially fractionate the mixed population of BchNB proteins over Ni^2+^-nitriloacetic acid (NTA) and Strep-Tactin-agarose resins. The final purified BchNB complex had one BchB subunit carrying a poly-histidine tag and the other a Strep-tag (Fig. 2e). The presence of the appropriate affinity tags over the sequential purification steps were confirmed by western blotting using anti-His and anti-Strep antibodies (Fig. 2g, i and Supplemental Fig. 1).

**Figure 2.**
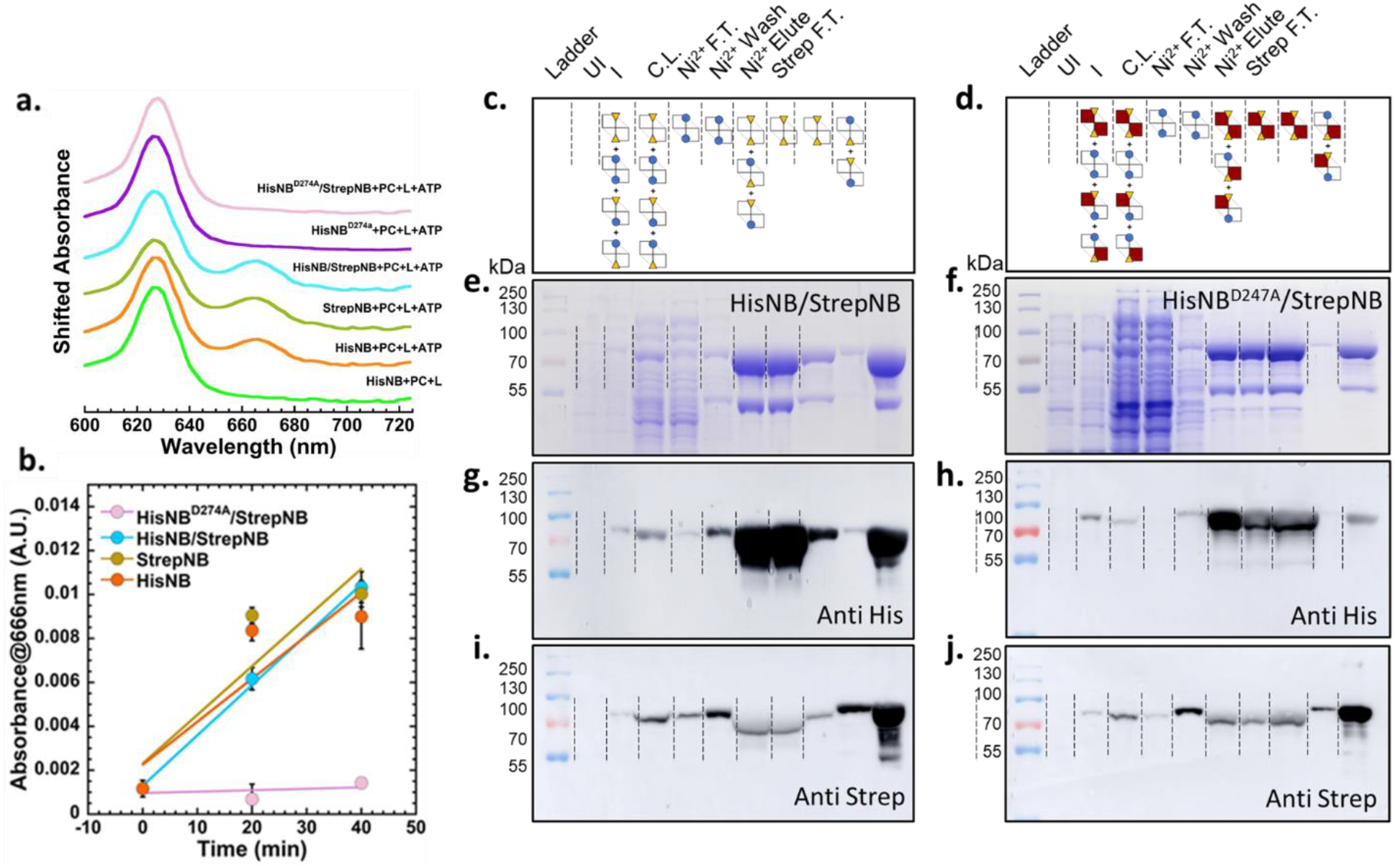
A D274A substitution within one BchB subunit in the tetrameric BchNB complex abolishes Pchlide reduction activity in DPOR. **a.** Pchlide reduction activity of wildtype, mutant, and half-mutant BchNB complexes were assayed by monitoring the absorbance of Pchlide and Chide upon addition of BchL and ATP to each reaction. Absorbance spectra of the reactions are shown, but shifted relative to each other to clearly showcase the traces. The individual reactions are respectively labeled above each trace. **b.** Quantitation of time course of Pchlide reduction. **c. & d.** Cartoon representation of predicted protein pools in each fractionation. Poly-His and Strep-tags are denoted as yellow triangles and blue circles respectively. BchN is not depicted. Wild-type BchB are shown as white squares and BchB^D274A^ is denoted as red squares. **e. & f.** Coommassie-stained SDS-PAGE analysis of the fractionated protein samples. **g-i.** Western blot analysis was used to identify and ascertain the purification of Poly-His and/or Strep-tagged heteromeric protein complexes. The final purified BchNB complex shows the presence of heteromeric complexes carrying both affinity tags. Thus, the purified proteins carry either wt-wt, or wt-mutant BchNB heteromeric complexes.

To generate a half-active BchNB complex, we introduced an Asp274 to Ala substitution only in the BchB subunit carrying the poly-histidine tag. Asp274 acts as one of the necessary proton donors for Pchlide reduction, the other coming from the Pchlide itself (Fig. 1c). Asp274 interacts with the Pchlide molecule and substitution of Asp274 to Ala has been shown to perturb substrate reduction in DPOR(3). When expressed along with wild-type BchN and wild-type Strep-tag BchB, a mixture of BchB complexes were generated (Fig. 2d) carrying either entirely wild-type BchNB (strep-B:[BchN]_2_:strep-B); mutant BchNB (his-B^D-A^:[BchN]_2_:his-B^D-A^); or half-active BchNB (his-B^D-A^:[BchN]_2_:strep-B). Using the dual-affinity purification strategy we were able to isolate the half-reactive BchNB (his-B^D-A^:[BchN]_2_:strep-B; Fig. 2f) and confirmed the presence of both affinity tags by western blotting using the above described antibodies (Fig. 2h, j and Supplemental Fig. 1).

We next monitored Pchlide reduction activity of the dual-tagged wild-type and half-active BchNB complexes by mixing them with Pchlide, BchL and ATP. While wild-type BchNB carrying either affinity tag reduces Pchlide (Fig. 2a, b), the half-active BchNB is defective for Pchlide reduction (Fig. 2a, b). These data show that two functional halves in the context of the α_2_β_2_ BchNB tetramer are required for substrate reduction.

The predicted sequence of Pchlide-reduction events in DPOR involve binding of Pchlide to BchNB, the binding of BchL to BchNB and the respective electron transfer events. In addition, ATP binding and hydrolysis within the BchL complex are also elemental steps in substrate reduction(16, 17). Since the half-active BchNB proteins were defective for overall Pchlide reduction, we next sought to identify where the enzyme stalled in the catalytic cycle. Thus, we focused on measuring the electronic properties of the [4Fe-4S] clusters of BchNB using electron paramagnetic resonance (EPR) spectroscopy. Unfortunately, the protein yields of the half-active BchNB protein were minimal after the dual-affinity purification and hence precluded us from performing detailed EPR studies. Hence, we investigated the spectral properties of wild type and variant (BchNB^D274A^) DPOR complexes to capture the steps in the electron transfer cycle.

### The [Fe-S] cluster of BchNB matches a simulated s=7/2 spin state which can be pre-reduced, and re-primed for sequential reactions - a ‘deficit spending’ model

The Asp274 residue in BchB is unique in its role as a proton donor to Pchlide(3). Structurally, it functions in *trans*, where Asp274 from one half of the BchNB tetramer serves as the proton donor to the Pchlide molecule bound to the active site of the opposing half BchNB (Fig. 1c)(3, 4). Thus, this residue serves as a key communication element between the two halves of the α_2_β_2_ BchNB tetramer. The [4Fe-4S] cluster of BchNB is ligated by three Cys residues from BchN (C29, C54 and C115) and one Asp residue from BchB (D36; Fig. 3a). To better understand the catalytic events in DPOR, we first focused on the intrinsic electron transfer properties of the [4Fe-4S] cluster of BchNB.

**Figure 3.**
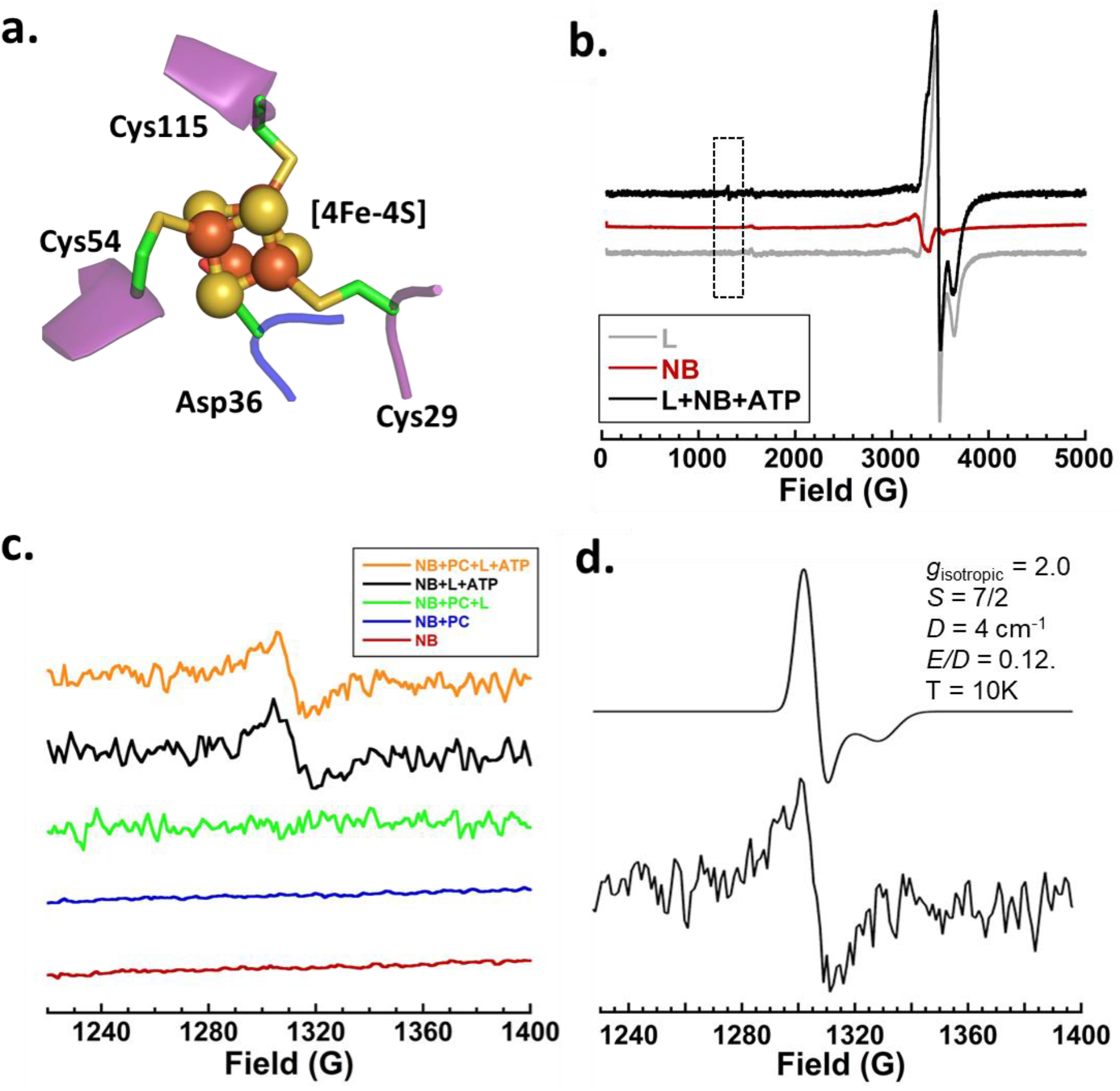
The [4Fe-4S] Cluster of BchNB is EPR active during Pchlide reduction and displays a novel S=7/2 signal. **a.** Structure of the [4Fe-4S] cluster of BchNB (generated from PDB ID:2YNM) shows coordination by three Cys residues from BchN and one Asp residue from BchB. **b.** Complete EPR spectra of BchL shows signals around ~3500 G, but no spectra in the ~1250 G range. BchNB shows a smaller EPR signal in the ~3400 G range that overlaps with BchL. When BchL and BchNB are mixed with ATP, a new (very small) signal arises in the ~1250 G region. **c.** EPR spectra recorded in the 1200-1400 G region show a lack of EPR signal for BchNB in the absence (red) or presence (blue) of Pchlide. No signal is observed when BchL, BchNB and Pchlide are incubated in the absence of ATP (green). A reaction containing BchL and BchNB in the presence of ATP, but lack of Pchlide, show electron transfer from BchL to BchNB (black). Thus, the component proteins can form a complex in the absence of Pchlide. When BchNB, BchL, ATP and Pchlide are present, the steady-state reaction shows the presence of the peak (orange). Thus, post reduction of Pchlide to Chlide, the [4Fe-4S] cluster of BchNB is in a pre-reduced state. **d.** Experimentally derived (bottom trace) and simulated (top trace) EPR spectra of a S=7/2 spin state of the [4Fe-4S] cluster of BchNB. Experimental and simulation parameters used for data fitting are denoted.

The EPR spectral properties of the BchNB [4Fe-4S] cluster have not been thoroughly characterized. Thus, we first measured the EPR spectra of BchNB alone, in complex with its substrate (Pchlide), and upon addition of BchL (+/− ATP). The [4Fe-4S] cluster of BchNB is EPR silent (Fig. 3b; red trace). In comparison, the [4Fe-4S] cluster of BchL produces a large signal in the ~3500 G range as reported previously (Fig. 3b; grey trace)(18). Even at 3-fold higher BchNB concentrations (90 μM tetramer) in the EPR reactions no EPR signature is observed (Fig. 3c; red trace). When Pchlide is added to BchNB, we do not observe a change in signal (Fig. 3c; blue trace). When BchNB, BchL and Pchlide are mixed together in the absence of ATP, the signal for the [4Fe-4S] cluster for BchL is observed, but no new signals for BchNB are observed (Fig. 3c; green) trace. This finding is not surprising as ATP binding to BchL is required to drive complex formation between BchNB and BchL.

When we introduce BchL and ATP to the reaction containing BchNB in the absence of Pchlide, a peak appears at ~1300 G (Fig. 3b, c; black traces). This suggests that ATP-bound BchL forms a complex with BchNB in the absence of Pchlide and donates an electron to the [4Fe-4S] cluster of BchNB. When the entire reaction is reconstituted with BchL, BchNB, Pchlide, and ATP, the steady state reaction results in the presence of the ~1300 G peak (Fig. 3c; orange trace). By modulating the temperature and reaction conditions, we were able to capture this EPR signal of the uniquely ligated BchNB cluster, with maximal signal observed at 10K (Supplemental Fig. 2 and Supplemental Note 1). The spectrum matches a simulated s=7/2 spin state (Fig. 3d). Previous models for DPOR suggest that the [4Fe-4S] cluster of BchNB exists in an oxidized state, binds to Pchlide, followed by complex formation with BchL and the transfer of the first electron from BchL to BchNB and then onto Pchlide. In our experiments, since BchL can reduce BchNB in the absence of Pchlide, we suggest an alternate possibility where BchNB could be pre-reduced in the cell followed by Pchlide binding, and the subsequent transfer of the first electron to Pchlide. This model is congruent with the “deficit spending” mechanism proposed for nitrogenase(15), wherein the first electron donation to substrate is accomplished before ATP-dependent reduction from the Fe-protein (electron donor component).

We next measured the kinetics of substrate reduction in DPOR by following the appearance of the absorbance signal at 666 nm (Fig. 4a), and the appearance of the ~1300 G EPR signal (Fig. 4b). Reduction reactions containing BchL, BchNB, and Pchlide were initiated by adding ATP and quenched samples were assessed for electron transfer (EPR) and Pchlide formation (absorbance). The appearance of the EPR signal correlates with the formation of Chlide, albeit at a slower rate (Fig. 4c), suggesting that a) the EPR signal represents electron transfer to the [4Fe-4S] cluster of BchNB. b) Since rate of appearance of the EPR spectra lag behind the Chlide formation (absorbance signal), the EPR signal likely reflects the re-reduced cluster on BchNB. Thus, during the reaction, the BchNB cluster is always in a reduced state, and primed to donate an electron to

**Figure 4.**
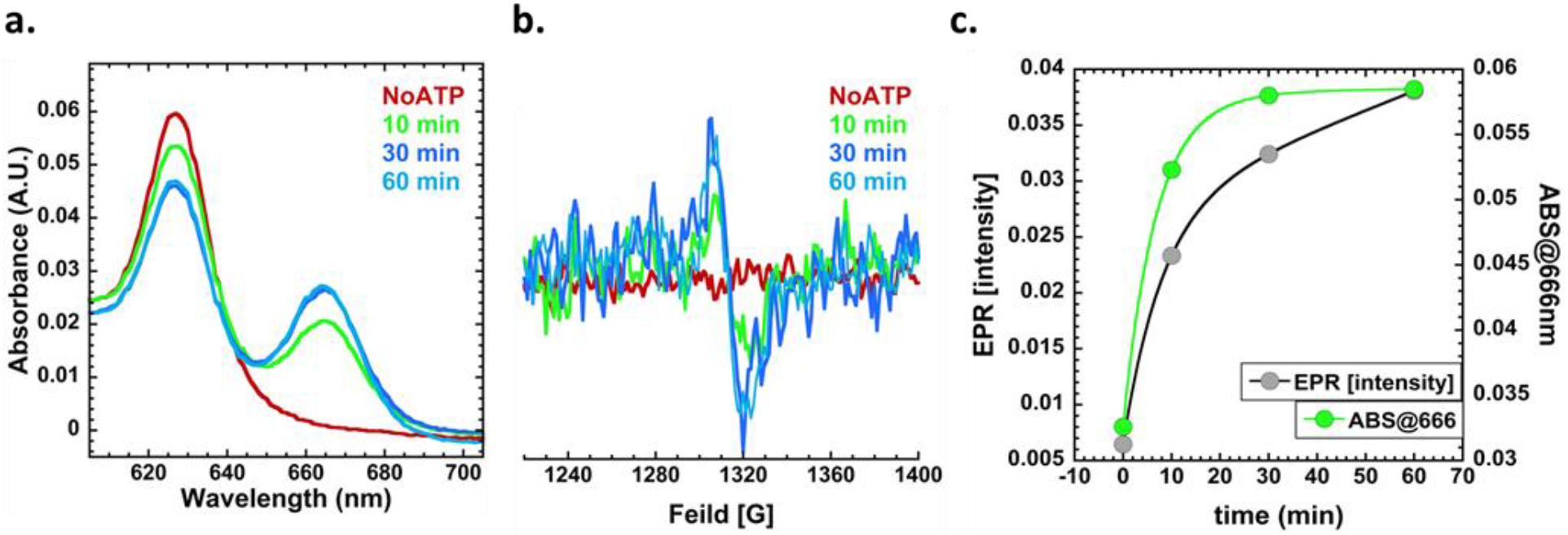
The [4Fe-4S] cluster of BchNB is in a reduced state after Pchlide reduction. BchL, BchNB, and Pchlide were mixed and the reaction was initiated by the addition of ATP. At the denoted times: **a.** Samples were removed and the amount of Pchlide reduced was spectroscopically determined after acetone extraction of the pigments. Chlide formation was detected as a function of time. **b.** EPR spectra was also measured for the samples and the intensity of the signal at ~1300 G increased as a function of time. **c.** EPR intensity at ~1300 G (grey circles, left axis) and absorbance at 666 nm (green circles, right axis) increased as a function of time. The kinetics of the Pchlide reduction signal reached maxima before the EPR intensity saturated suggesting that the [4Fe-4S] cluster of BchNB is re-reduced post Pchlide reduction, likely priming it for the deficit-spending mechanism for the next round of electron transfer.

Pchlide during the next substrate reduction cycle - as would be expected in the ‘deficit spending’ model.

### Asp274 to Ala substitution reveals unresolved electron transfer intermediates in BchNB and a sequential substrate reduction mechanism within the two halves

Establishment of the order of Pchlide binding to BchNB is necessary to understand the mechanism of ET. From the half-reactive BchNB experiments we deciphered that multiple rounds of ET do not occur when one or both BchNB halves carry a substitution in the proton donor residue - Asp274. Thus, we proposed that a stalled BchNB intermediate could be identified through EPR for the BchNB^D274A^ complex. Interestingly, the EPR signature of BchNB^D274A^ (in the absence of Pchlide, BchL or ATP) showed the presence of a peak at ~1300 G, indicative of the presence of a reduced [4Fe-4S] cluster (Fig. 5a). This is in stark contrast to wild type BchNB that is EPR silent (Fig. 5a). This suggests that the substitution at Asp274 influences the electronic properties of the [4Fe-4S] cluster either by physically promoting structural changes in the protein that make the cluster more accessible to the reductant (dithionite) in the reactions, or by propagating/perturbing a network of amino acid interactions that change the reduction potential of the cluster and making it EPR visible.

**Figure 5.**
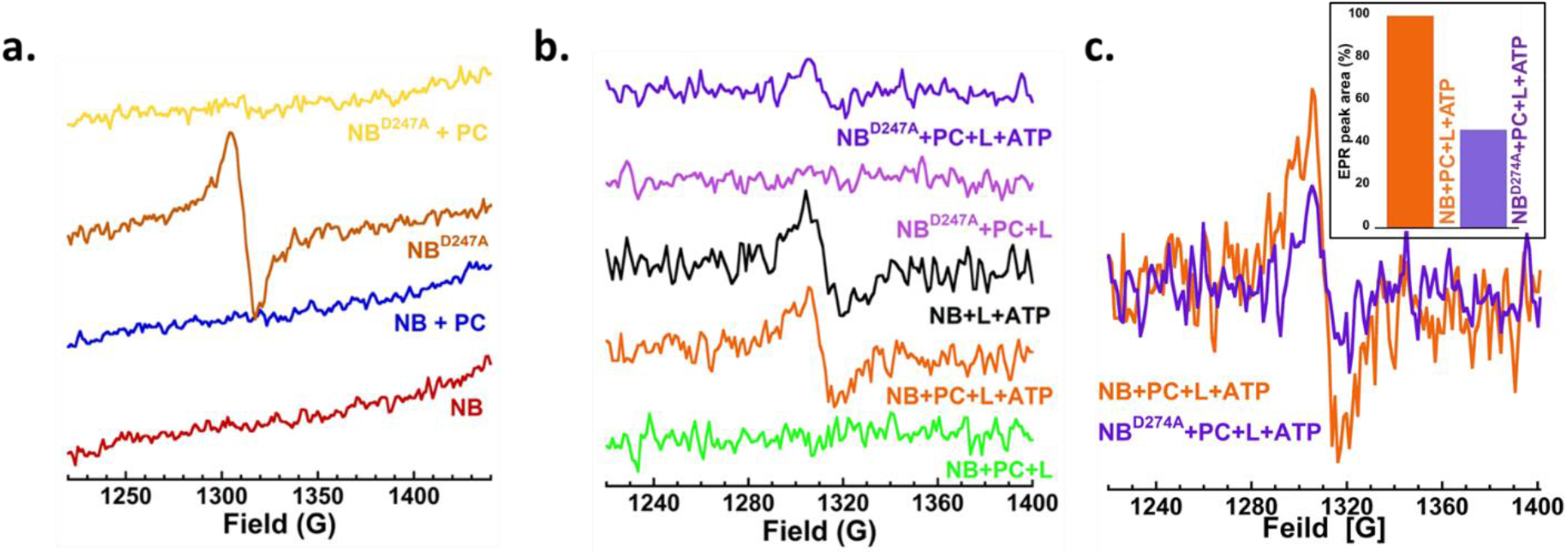
BchNB^D274A^ is synthesized with electrons pre-loaded and reveals a deficit-spending like mechanism for the first electron transfer event. **a.** Comparison of EPR spectra of wild type BchNB (red) versus BchNB^D274A^ (orange) reveals a pre-reduced [4Fe-4S] cluster in BchNB^D274A^. When Pchlide is added, the electron is donated and cluster becomes EPR-silent (yellow). **b.** EPR spectra of wild type BchNB and BchNB^D274A^ in the presence and absence of BchL and ATP reveal electron transfer properties of the DPOR complex. No ET is observed between BchL and BchNB in the absence of ATP, but presence of Pchlide (green). ET from BchL to BchNB is observed in the presence of ATP, and in the absence of Pchlide (black). Steady-state ET is observed when all components of the reaction are present (orange). The preloaded electron in BchNB^D274A^ is transferred to Pchlide in the presence of BchL, and in the absence of ATP (purple), thus supporting evidence for a deficit-spending mechanism. When all the reaction components are present with BchNB^D27A^ (blue), the EPR intensity is half that observed for wild type BchNB. **c.** Overlay of the EPR spectra for wild-type BchNB (orange) and BchNB^D274A^ (blue) in the presence of BchL, Pchlide, and ATP, show that the amplitude of the BchNB^D274A^ intensity is ~50% of the wild-type reaction (insert). Thus, BchNB and BchL form a ATP-dependent complex in the absence of Pchlide and can donate an electron from BchL to BchNB and then onto Pchlide, followed be reduction of the BchNB [4Fe-4S] cluster

Analysis of the crystal structures of the Pchlide-free(19) and Pchlide-bound BchNB(3) crystal structures show large scale conformational changes that might likely explain this difference (Supplemental Fig. 3). The collapse of the binding pocket in wild type BchNB, in the absence of Pchlide, also likely serves as a protective mechanism for the [4Fe-4S] cluster. We recently reported a unique protective mechanism in the BchL protein where a flexible disordered region in the N-terminus bound across the [4Fe-4S] cluster and autoinhibited BchL activity(20). The inhibition is released through conformational changes upon ATP binding. We propose that the collapse of the Pchlide binding pocket might serve a similar protective role in BchNB as the cluster is not able to be reduced. Binding of Pchlide, or BchL (or both), likely triggers the necessary conformational changes and primes the [4Fe-4S] cluster for accepting electrons. BchNB can bind to Pchlide in the absence of BchL; similarly, BchL can bind to BchNB in the absence of Pchlide and donate an electron (in the presence of ATP). Thus, Pchlide or BchL binding to BchNB are likely not mutually exclusive events and a specific order of binding may not dictate overall substrate reduction activity.

Since BchNB^D274A^ is defective for Pchlide reduction, but is preloaded with electrons, we next tested whether ET occurs. When incubated with Pchlide, the EPR spectra for the [4Fe-4S] cluster of BchNB^D274A^ disappears (Fig. 5a). Thus, BchNB^D274A^ donates its pre-loaded electrons to Pchlide. The complete disappearance of the EPR signal implies that the first ET event occurs in both halves. Interestingly, when BchNB^D274A^ is incubated with BchL, Pchlide and ATP, the EPR signal at ~1300 G is resurrected (Fig. 5b). However, the intensity of the EPR peak is almost exactly 50% of the amplitude of the signal observed for wild-type BchNB under similar conditions (Fig. 5c). This is indicative of a stalled intermediate where re-reduction of only one of the two [4Fe-4S] clusters of BchNB has occurred. The data also affirm our interpretation that the EPR signal at ~1300 G for wild type BchNB (and the BchNB^D274A^ without Pchlide) reflects two total electrons per tetramer - one per BchNB half.

These findings further support the sequential ET model in DPOR where events in one half control activity in the other. If the two halves were to act independently, a) the half-reactive DPOR complex should have been able to retain partial Pchlide reduction activity, but this is not the case (Fig. 2. b) Similarly, in the EPR analysis, for an independent model, the amplitude of the spectra for BchNB^D274A^ should have been either around 100% or 0% of wild-type BchNB, but this again is not the case (Fig. 5c). Thus, we propose that an intrinsic functional asymmetry exists in DPOR.

### Direct measurements of Pchlide binding to BchNB reveal a substrate sensing mechanism that establishes functional asymmetry

Since the two halves in the BchNB complex appear to transfer electrons to Pchlide in a sequential manner, we next tested how asymmetry is established. Either BchNB, when synthesized, is intrinsically endowed with asymmetric Pchlide binding properties through differences in active-site conformations between the two halves. Alternatively, Pchlide binding to either active site is stochastic and changes in the active site upon Pchlide binding then sets asymmetry and allosterically controls Pchlide binding to the other half. To test these models, we directly monitored Pchlide binding by capturing the changes in fluorescence upon binding to BchNB.

A scan of the fluorescence properties of Pchlide shows an excitation and emission maxima at 440 and 636 nm, respectively (Supplemental Figure 4). Based on the spectral features, we excited a sample of Pchlide at 440 nm and measured the change in fluorescence (Fig. 6a). When BchNB is added to the reaction, the fluorescence emission of Pchlide increases ~750-fold (Fig. 6a). The utility of Pchlide fluorescence to investigate its binding to other proteins has been controversial. One such study using protochlorophyllide reductase (POR) showed no significant changes in Pchlide fluorescence upon binding to Pchlide; as non-specific binding to BSA also generated similar changes in fluorescence(21). Thus, such changes in Pchlide fluorescence were ascribed to non-specific solution partition effects. This is not the case for Pchlide binding to the BchNB complex. Under our reaction conditions, BSA does not produce a change in Pchlide fluorescence (Fig. 6a). Thus, for BchNB, changes in Pchlide fluorescence are an excellent measure of binding.

**Figure 6.**
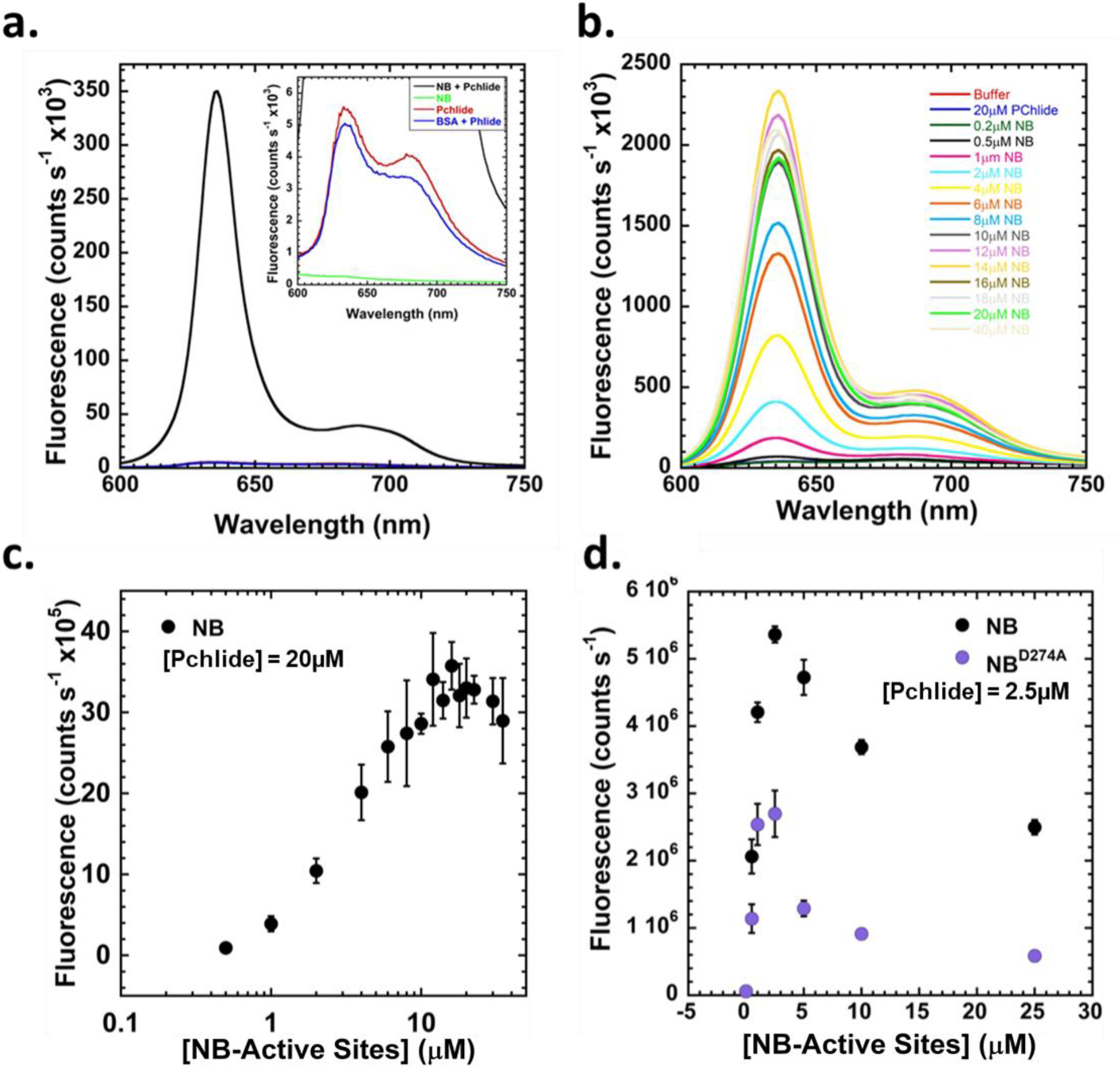
BchNB senses Pchlide and conformationally induces its fluorescence enhancement. **a.** Fluorescence of Pchlide was measured in the absence or presence of BchNB. BchNB binding to Pchlide generates a ~700-fold increase in Pchlide fluorescence. BSA does not change the fluorescence properties of Pchlide (insert). **b.** Changes in fluorescence of Pchlide (20 μM) was measured as a function of increasing concentration of BchNB (concentration denoted in terms of the number of Pchlide binding sites), and **c.** the change in fluorescence was plotted as a function of increasing BchNB concentration. Pchlide fluorescence saturates when stoichiometric number of binding sites on BchNB are available. **d.** Pchlide (2.5 μM) was mixed with increasing concentrations of wild type BchNB or BchNB^D274A^, and the change in fluorescence was measured. The fluorescence maxima is obtained when stoichiometric amounts of Wild type BchNB and BchNB^D274A^ Pchlide binding sites are available. However, the amplitude of fluorescence for BchNB^D274A^ is half as observed for wild type BchNB. When excess BchNB is added to the reaction, the complexes shift from two Pchlide bound per BchNB or BchNB^D274A^ to one Pchlide bound per complex.

Since the fluorescence signal arises from Pchlide, we performed binding experiments with a fixed concentration of Pchlide and titrated in increasing concentrations of the BchNB tetramer (Fig. 6b). An increase in Pchlide fluorescence is observed as of function of BchNB concentration (Fig. 6c). The signal plateaus when enough BchNB is present to bind all the Pchlide molecules in the reaction. Curiously, when an excess of BchNB is added to the reaction the fluorescence does not plateau. Instead, a drop in the Pchlide fluorescence is observed (Fig. 6c). As the concentration of BchNB is increased in the reaction, the equilibrium likely shifts from a 2-Pchlide bound BchNB complex (higher fluorescence) to a 1-Pchlide bound complex (lower fluorescence). The quantum yield of the 2-Pchlide bound complex is likely different than the 1-Pchlide bound complex. Given the asymmetry in BchNB activity, we propose that one Pchlide is bound in a conformation different than the other. The conformational differences likely contribute to two different quantum yields for Pchlide fluorescence when bound to BchNB.

When similar binding experiments are performed with the BchNB^D274A^ variant, we observe a similar overall profile where an increase in Pchlide fluorescence is observed. The signal saturates stoichiometrically, as observed for wild-type BchNB. Thus, both active sites are Pchlide bound (Fig. 6d). However, for BchNB^D274A^, the fluorescence quantum yield upon reaching stoichiometry is half that observed for wild-type BchNB. These data suggest that either the conformational positioning or electronic landscape of the bound Pchlide molecules are different between BchNB^D274A^ and wild-type BchNB.

Extrapolating back to the EPR results, the spectra obtained for the wild type BchNB likely reflect one electron transferred to each Pchlide molecule. This could function as a sensing step to recognize and lock in the Pchlide within the active site. The EPR spectra for BchNB^D274A^, measured under the same conditions, also show ET to the bound Pchlide molecules (Fig. 5a). In BchNB^D274A^, the inability to donate the first proton from Asp274 to Pchlide is perturbed.

In the presence of BchL and ATP, complex formation between Pchlide-bound-BchNB and ATP-bound-BchL occurs. This re-reduces the [4Fe-4S] cluster on BchNB. The extent of re-reduction is 50% in the BchNB^D274A^ variant and thus Pchlide reduction is not achieved in either active site. This evidence points to sequential ET in the DPOR system and the Asp274 residue serves as a key sensor in communicating the electron transfer status between the two halves.

## Discussion

The oligomeric α_2_β_2_ structural arrangement observed in the BchNB complex of DPOR is found in a host of enzymes, especially in enzymes that catalyze ET reactions. These enzymes have two diametrically situated, identical active sites. In DPOR, extensive structural contacts between the two active sites are a prominent feature with one half contributing a key proton donor (Asp274) in *trans* to the Pchlide molecule bound in the active site of the opposing subunit(3, 4). Thus, allosteric and direct communication between the two halves likely controls the catalytic steps in the ET and Pchlide reduction mechanisms. Our findings support this model and show communication between the two active sites. In the half-reactive engineered version of BchNB, where one site is wild-type and other carries a D274A substitution, Pchlide reduction activity in both halves is abolished. Thus, in this scenario, the half-reactive enzyme does not go through multiple rounds of ET required for Pchlide reduction. The proposed sequence of catalytic events for DPOR activity involves:

1. chlide binding to both active sites of BchNB. Either BchNB, when synthesized in the cell, is structurally asymmetric with respect to the organization of the two identical active sites. This could dictate the order of Pchlide binding to the two active sites. Alternatively, the active sites are identically poised to accept Pchlide and stochastic binding of Pchlide to one or the other site establishes asymmetry within BchNB. Contrary to previous models, our data show that the [4Fe-4S] cluster of BchNB exists in a reduced state and is able to donate the first electron to Pchlide upon binding in the absence of BchL. In the steady state experiments, the EPR spectra corresponding to the [4Fe-4S] cluster of BchNB shows a persistent reduced state well after the time course of Pchlide reduction (Fig. 4). This mechanism is similar to the ‘deficit spending’ model proposed for nitrogenase, where the first ET in the MoFe-protein occurs before binding of the Fe-protein(15). Whether this first ET occurs sequentially in both active sites of BchNB, or just within one site (thus maintaining asymmetry) is yet to be determined. Such sequential ET occurs in nitrogenase where the ET in one half allosterically suppresses catalytic events in the other(22, 23). Another interesting observation is the difference in pre-reduced states of our wild-type BchNB and the BchNB^D274A^ variant. The [4Fe-4S] cluster of BchNB^D274A^ is always pre-reduced and is visible in EPR (Fig. 5a). The wild-type BchNB protein is EPR silent (Fig. 3c). This difference suggests conformational differences around the [4Fe-4S] cluster between the wild-type and variant BchNB complexes. However, both proteins bind to two Pchlide molecules (Fig. 6d). We propose that the Asp276 residue is used to sense and communicate the presence of Pchlide molecule within the active site and the deficit spending mechanism might be an integral part of this process. This sensing and selection of Pchlide likely serves two roles - a) it correctly positions the porphyrin-ring structure of Pchlide within the active site, and b) it helps DPOR differentiate between Pchlide and Chlide. Chlide is the reduced product of Pchlide and must dissociate from the active site of DPOR. Chlide then binds to the active site of chlorophyllide oxidoreductase (COR), the next enzyme in the pathway. COR is proposed to be structurally similar to DPOR, but the difference between Pchlide and Chlide is one double bond in the C17=C18 position of the porphyrin ring. Thus, the initial positioning and sensing of Pchlide by the Asp274 residue likely serves as a ‘substrate-check’ in the active site of BchNB.
2. The next step in the DPOR mechanism is the binding of BchL to BchNB. We do note here that while its logical to think of Pchlide binding to BchNB as the first step, followed by complex formation with BchL, these steps need not be sequential or dependent on one other. BchL forms a complex with BchNB in the absence of Pchlide and transfers electrons (Fig. 3c). Thus, a direct measure of the kinetics and thermodynamics of complex formation between BchNB and BchL, and the influence of Pchlide, would be required to better understand the complexities underlying complex formation in DPOR. Similarly, when BchL binds to BchNB, these events can occur on both halves of the BchNB complex. Whether these binding events are stochastic, cooperative/independent, and how they contribute to overall asymmetry remains to be investigated. The crystal structure of DPOR bound to the ATP-analog ADP-AlF_3_ stabilizes a transition-state complex with BchNB bound to BchL on both halves(4). In the absence of ATP, there is no ET from BchL to BchNB (Fig. 3c). Thus, at a minimum, complex formation between BchL and BchNB is transient and coupled to ATP binding and hydrolysis within BchL. We recently showed that a disordered region in the N-terminus of BchL is auto-inhibitory and ATP-binding relieves the inhibition to drive complex formation with BchNB(20). This regulation appears unique to DPOR as neither the Fe-protein of nitrogenase or the BchX protein of COR possess this regulatory N-terminal region.
3. ET from the [4Fe-4S] cluster of BchL to the [4Fe-4S] cluster on BchNB and then onto Pchlide occurs upon complex formation. As stated above, whether ET occurs independently within the two halves remains to be established. Our data provides strong evidence that communication regarding ET from one half is relayed to the other through Asp274. Substitution of Asp274 in just one half of BchNB stalls Pchlide reduction. Thus, the two halves are synchronized with respect to the steps in their Pchlide reduction cycles. In nitrogenase, we showed negative cooperativity and allosteric communication between the two halves(22). ET occurs first in one half, and this process is suppressed in the other, thus establishing a sequential ET mechanism between the two halves. Since the half-active DPOR complex fails to reduce Pchlide in both halves, we propose that such asymmetry also exists in the DPOR system.

The commonalities between DPOR and nitrogenase, with respect to communication and allostery between the two halves suggest that sequential ET might be a theme embedded in other such α_2_β_2_ structured enzymes. Evolutionary functional advantages to such structures could be used to direct the flow of electrons, substrate-binding selectivity, and overall efficiency of ET. Both nitrogenase and DPOR catalyze substrate reduction reactions requiring multiple rounds of ET. The allosteric and sequential ET process could also be utilized to count/calibrate the number of electrons accumulate at the metal clusters and on the substrate. It would be interesting to explore such mechanistic differences between α_2_β_2_ enzyme complexes that catalyze single versus multiple ET dependent reduction chemistries.

## Materials and Methods

### Generation of protein expression constructs

Plasmids used to recombinantly produce BchL and BchNB are described. The appropriate N-or C-terminal poly-histidine or strep-tags were engineered onto BchB using Q5 site-directed mutagenesis (New England Biolabs, Ipswich, MA). The primers used to generate the corresponding sequence change are listed (Supplemental Table 1). The amino acid sequences of the encoded proteins are also provided (Supplemental Information).

### Generation of Pchlide

Pchlide was generated as described(24) using a *Rhodobacter sphaeroides* ZY-5 strain harboring a deletion of the BchL gene (a kind gift from Dr. Carl Bauer, Indiana University)(8).

### Protein expression and purification

Wild-type/mutant BchNB and BchL protein complexes were purified as described(24). The following modified procedure was used for purification of the half-reactive BchNB complex. Plasmids encoding BchN, His-BchB and Step-BchB were co-transformed into BL21(DE3) SufFeScient cells. The transformants were grown, and protein overproduction was induced as described for the wild-type BchNB complex. After lysis (as described), the clarified lysates from His-tagged or dual tagged constructs were loaded onto an Nickel-nitrilotriacetic acid (Ni^2+^-NTA; 1.6 mL suspended beads/L growth) column (Thermo Scientific) equilibrated with 10 column volumes (CVs) STD (100 mM HEPES PH7.5, 150 mM NaCl) buffer. Non-specifically bound proteins were washed off with 10 CVs STD buffer containing 20 mM imidazole. Bound proteins were eluted into a septum sealed bottle using 30 mL STD buffer containing 250 mM Imidazole. The clarified lysates from Strep-tagged BchNB were loaded onto a Strep-Tactin Sepharose column (IBA, Germany; 1.6mL resuspended beads/L growth) equilibrated with 10 CVs STD buffer. Post washing with 10 CVs STD buffer, bound proteins were eluted using 30 mL STD buffer containing 25 mM Desthiobiotin (IBA). Asymmetric or dual tagged BchNB constructs followed the methods described for Ni^2+^-NTA columns and for Strep columns but performed sequentially. Eluted BchNB proteins were concentrated using a spin concentrator (30 kDa molecular weight cut-off). Proteins were aliquoted in the glove box into 1.2 mL cryo-tubes which have a gasket sealed cap (Cat. #430487 Corning). Closed tubes with protein were removed from the glove box and flash frozen using liquid nitrogen and stored under liquid nitrogen. Protein concentrations were determined using Bradford reagent with bovine serum albumin as reference.

### Western Blotting

Western blots were performed on samples after separation on 10% SDS-PAGE. Proteins were transferred (100 mA for 90 min) on to nitrocellulose membranes. Membranes were washed thrice with TBST (Tris buffered saline plus 0.1 % (v/v) Tween-20), and rocked for 15 minutes after the third rinse. Membranes were blocked with 5 % (w/v) milk in TBST for 1 hour at 25 °C. Membranes were then washed with TBST thrice as before, then exposed to primary antibody. Membranes were exposed to StrepMAB-Classic HRP conjugate (IBA) primary antibody (1:34000 dilution in TBST) for 1 hour while rocking at 25 °C and/or anti-poly-histidine antibody, HRP conjugate (Invitrogen) (1:1000 dilution in TBST plus 1 % (w/v) milk) overnight (~17 hours) while rocking at 4 °C. Membranes were then washed, rocked for 3 hours after the third rinse before adding detection reagent (Pierce Fast Western Blot Kit, ECL substrate, Thermo Scientific). Blots were imaged using an AI 600 imager (GE Healthcare).

### Assay for substrate reduction by DPOR

Reduction of Pchlide to Chlide was measured spectroscopically by mixing BchN-BchB protein (3 μM tetramer), BchL-protein (9 μM dimer), and 35 μM Pchlide, in the absence or presence of ATP (3 mM) in STD buffer containing 10 mM MgCl_2_. 40 μl of these reactions were quenched with 160 μl of 100 % acetone (80% v/v final concentration). The acetone extraction was then spun down in a table-top centrifuge at 13,226 g for 4 minutes to pellet precipitated protein components. 160 μl of the supernatant was transferred to a cyclic olefin half-area well plate (catalog #4680 Corning) and absorbance scans from 600 nm to 725 nm were recorded on a SpectraMax i3x plate reader (Molecular Devices). Chlide appearance was represented as the absorbance value at 666nm.

### EPR Spectroscopy

EPR spectra were obtained at 10 K on an updated Bruker EMX-AA-TDU/L spectrometer equipped with an ER4112-SHQ resonator (9.48 GHz) and an HP 5350B microwave counter for precise frequency measurement. Temperature was maintained with a ColdEdge/Bruker Stinger S5-L recirculating helium refrigerator, and an Oxford ESR900 cryostat and MercuryITC temperature controller. Spectra were recorded with either 0.3 G (3 □ 10-5 T) or 1.2 G (0.12 mT) digital field resolution with equal values for the conversion time and the time constant, 5.2 mW incident microwave power, and 12 G (1.2 mT) magnetic field modulation at 100 kHz. EPR simulations were carried out using Easyspin(25). Samples for EPR Spectroscopy. Reactions contained various combinations of BchL, BchNB, Pchlide, and ATP. 200 μl EPR samples contained 1.7 mM dithionite, and, where indicated in the figure legends, 40 μM BchL, 20 μM BchNB, 40 μM Pchlide, and 3 mM ATP. Samples from Fig. 5a contained 60μM BchNB, and/or 120μM Pchlide. All samples were incubated in the glove box for 60 minutes unless otherwise denoted. Protein samples were prepared and transferred to the EPR tubes in the glove box and stoppered with a butyl rubber stopper. Samples were removed from the glove box and immediately flash frozen in liquid nitrogen and then analyzed by EPR.

### Steady State Pchlide Binding Fluorescence titrations

2.0 mL mixtures of BchNB (various concentrations) and Pchlide (20 μM) were made in the glove box in STD buffer with 10 mM MgCl_2_, then incubated for 20 minutes before being transferred to an airtight 2 mL SEPTA screw cap cuvette (Firefly Scientific). Fluorescence emission spectra were recorded on a PTI fluorometer (Photon Technology International) excited at 440 nm, with excitation and emission slit widths of 1.5 mm and 3.0 mm respectively. PMT voltage and slit widths were maintained between replicates.

## Acknowledgements

This work was supported by a grant from the Department of Energy, Office of Science, Basic Energy Sciences (DE-SC0017866) to E.A. EPR was supported by an NSF Major Research Instrumentation award (CHE-1532168) to B.B. and by Bruker Biospin. E.I.C. was supported by a GAANN fellowship from the department of education and the Arthur J. Schmitt fellowship from Marquette University.

## Author Contributions

E.I.C. and B.B. performed experiments. E.I.C., B.B. and E.A. designed experiments and performed data analysis. E.A., B.B., and E.I.C. wrote the manuscript.

## Abbreviations

ET: electron transfer
DPOR: dark-operative protochlorophyllide oxidoreductase
Pchlide: protochlorophyllide
Chlide: chlorophyllide

