## Supplemental Information for "Substrate sensing institutes sequential and asymmetric electron transfer in the nitrogenase-like DPOR complex"

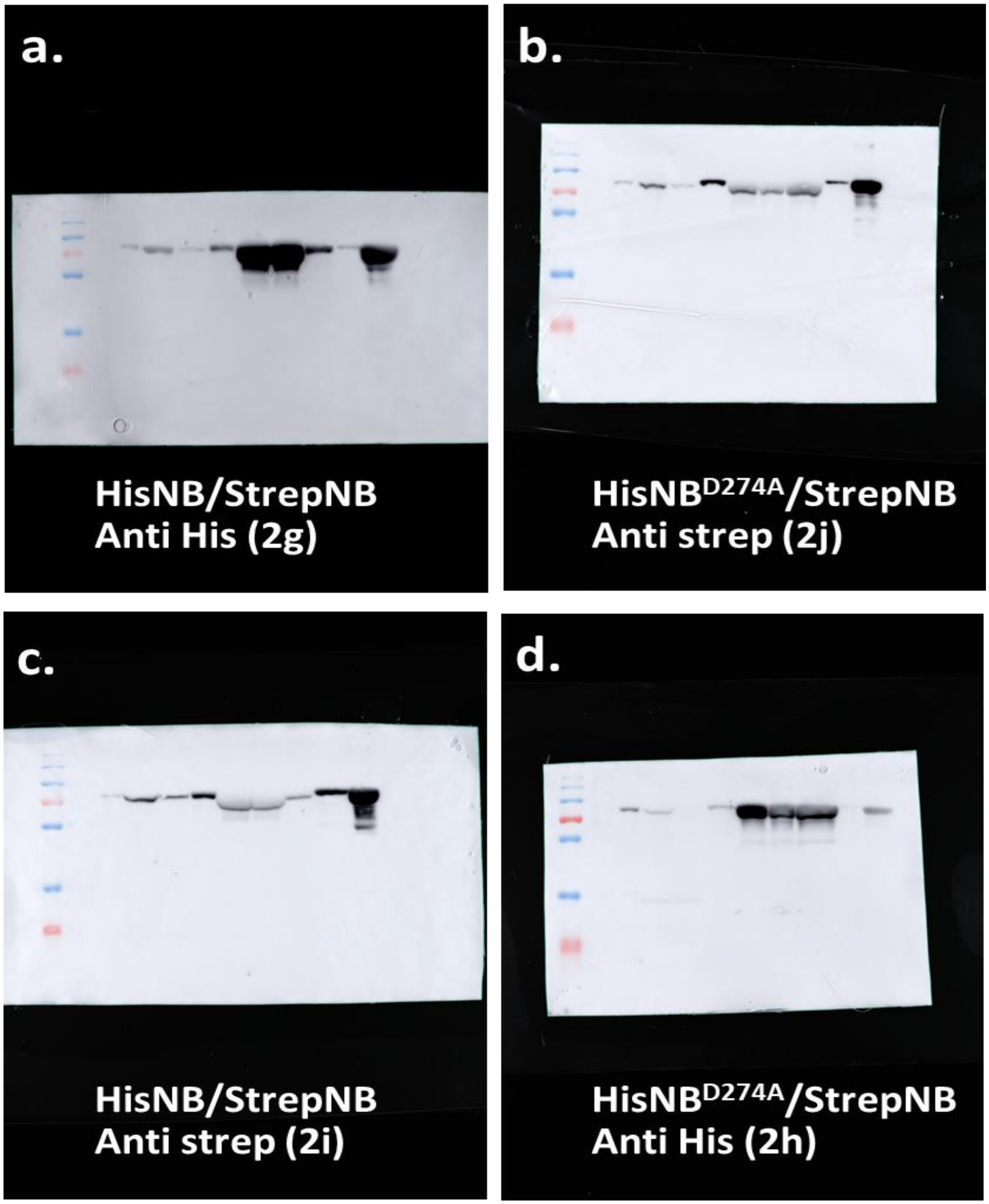


**Supplemental Figure 1. Raw images of unedited western blots of BchNB used in Figure .**
Western Blots used to generate data panels in Figures 2g., h., i., and j., respectively.

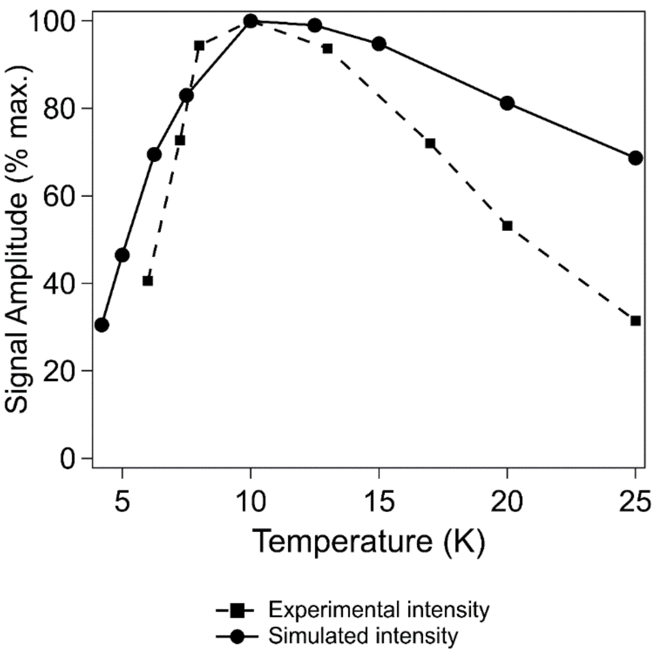


**Supplemental Figure 2. Simulated and experimental changes in EPR signal amplitude.** The signal intensity of the EPR spectral peak at G~1310 was integrated at the temperatures denoted above. Signals are plotted as a function of temperature relative to the maximal observed signal intensity at 10K. Simulated intensities are also plotted using the following parameters: $g_{isotropic}$ = 2.0; S = 7/2; D = 4 $cm^{-1}$; D = 0.12

**Supplemental Note 1: EPR characterization of the [4Fe-4S] cluster of BchNB.**
There has been very little characterization of the [4Fe-4S] cluster of BchNB by magnetic techniques; no MCD, Mössbauer, ENDOR etc. data have been reported. The only EPR signal that has been observed is the small, isotropic resonance at $g' \approx 5.15$ with a narrow window of temperatures in which it is detectable(1). Here, we have analyzed the EPR spectra of this cluster as a function of temperature (Supplemental Figure 2) and the data fit to a S=7/2 spin system. One potential concern is the very low intensity of this signal if it is proposed to be representative of a significant proportion of the clusters in the sample. The sharp drop in intensity at temperatures below about 7 K suggests that the low intensity is not simply due to very fast relaxation of the $S$ = 7/2 spin system. One possible explanation is that the $S$ = 7/2 spin state is actually the $S'$ = 7/2 *effective* spin state in a spin ladder(2). The theoretically possible spin states of a $[4Fe\text{-}4S]^{+}$ cluster are 1/2, 3/2,... 17/2. Depending on the overall zero-field splitting, which will in turn depend on the couplings between individual iron ions and pairs of ions in the cluster (Hagen 1985), either the highest or lowest spin state may represent a fully-occupied ground state (overwhelmingly, the $S$ = 1/2 state is observed for $Cys_4$-ligated $[4Fe\text{-}4S]^{+}$ clusters), or else the populations of the various $S'$ = 1/2, 3/2,... 17/2 effective spin states will occur in a temperature-dependent manner. Concerning relaxation, the higher spin states are expected to relax more rapidly such that the lower spin states will undergo saturation or rapid passage at low temperatures while the higher

spin states will only be observable at the lowest temperatures, if at all. We therefore speculate that the narrow temperature window for observation of this signal is due to temperature-dependent population of the $S'$ = 7/2 state. The value for $E/D$ (= 0.12) is particularly important here. The **g'**-tensor for $S$ = 7/2 is almost perfectly isotropic for this value, maximizing the *amplitude* of this signal, whereas for the other spin states the signal is highly rhombic and the EPR absorption is distributed over a wide field envelope and thus has very low intensity at any particular resonant field. Therefore, while each of the other states is paramagnetic, the $S'$ = 7/2 state may be the only EPR-detectable one at biological concentrations and where the spin density is distributed among the nine $S'$ spin manifolds at most temperatures. The observation that no spin states lower than $S'$ = 7/2 are observed may be an indication that the ground state is the lowest $S'$ = 1/2 state, as in other $[4Fe\text{-}4S]^+$ clusters, but that the zero-field splitting is much smaller.

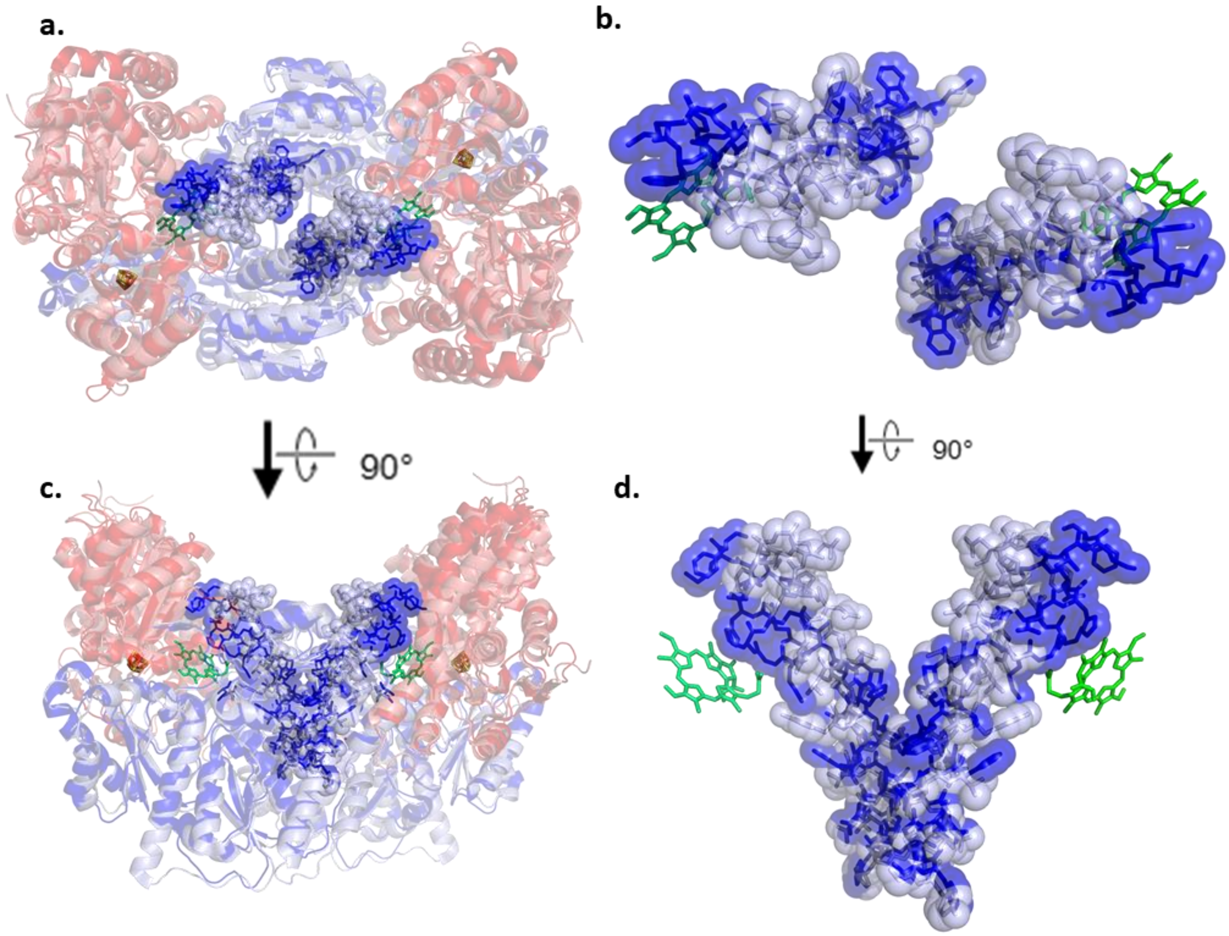


**Supplemental Figure 3. Conformational changes in BchNB upon binding to Pchlide.**
**a.** Top view of crystal structure of BchNB (dark red and dark blue, PDBID: 2XDQ) and Pchlide-bound BchNB (light red and light blue, PDBID: 3AEK) after structural alignment, shown as semi-transparent cartoon(s). [4Fe-4S] clusters and Asp274 (*R. sphaeroides* numbering) are shown as sticks, and Pchlide is shown as green sticks. The last alpha helix of either structure, which resides adjacent to Pchlide is shown as semi-transparent spheres, and opaque sticks. Color schemes are maintained throughout the entire figure. **b.** Highlights the terminal helix, and shows the relative movement upon Pchlide binding. **c.** Side view of the structure shown in a. **d.** Side view of the structure shown in b.

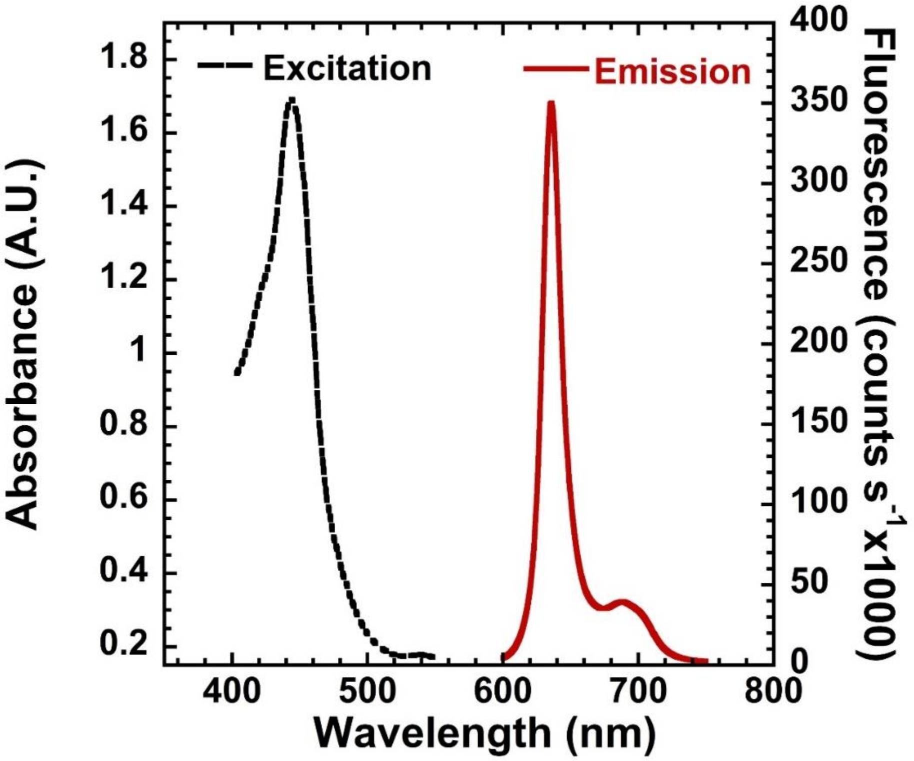


**Supplemental Figure 4. Fluorescence and absorbance properties of Pchlide- bound to BchNB.** Excitation-emission spectra of Pchlide (20 μM) bound to BchNB (10 μM)was measured by either following the changes in absorbance (left axis) or fluorescence (right axis). The peak excitation and emission wavelength was determined from this analysis.
